# Quantification of guanosine tetraphosphate and other nucleotides in plants and algae using stable isotope-labelled internal standards

**DOI:** 10.1101/2019.12.13.875492

**Authors:** Julia Bartoli, Sylvie Citerne, Gregory Mouille, Emmanuelle Bouveret, Ben Field

## Abstract

Guanosine tetraphosphate (G4P) and guanosine pentaphosphate (G5P) are signalling nucleotides found in bacteria and photosynthetic eukaryotes that are implicated in a wide-range of processes including stress acclimation, developmental transitions and growth control. Measurements of G4P/G5P levels are essential for studying the diverse roles of these nucleotides. However, G4P/G5P quantification is particularly challenging in plants and algae due to lower cellular concentrations, compartmentation and high metabolic complexity. Despite recent advances the speed and accuracy of G4P quantification in plants and algae can still be improved. Here, we report a new approach for rapid and accurate G4P quantification, applicable to plants and algae, which relies on the use of synthesised stable isotope-labelled as internal standards. We anticipate that this approach will accelerate research into the function of G4P signaling in plants, algae and other organisms.

## Introduction

Guanosine tetraphosphate (G4P) and guanosine pentaphosphate (G5P) are signalling nucleotides found in bacteria and photosynthetic eukaryotes that are implicated in a wide-range of processes including stress acclimation, differentiation and growth control (Field, 2018; Ronneau and Hallez, 2019). The accurate quantification of G4P and G5P is critical for studying the role of these signalling nucleotides. However, extraction and quantification of G4P/G5P can be challenging due to the vulnerability of these nucleotides to enzymatic and chemical degradation, their low basal concentrations, and high polarity. In bacteria G4P and G5P are routinely measured by radiographic detection on TLC plates after extraction from cells that have been globally labelled with P32, by growing them for several generations in the presence of 32P-labelled pyrophosphate. Ion exchange HPLC methods with UV detection have also been developed for the quantification of G4P and G5P in bacteria, and these methods have the advantage of allowing the simultaneous quantification of other nucleotides (Buckstein et al., 2008; Varik et al., 2017). More recently, a method using hydrophilic-interaction chromatography (HILIC) with mass spectrometry (MS) based detection and identification was also reported for bacteria (Zborníková et al., 2019). The quantification of other nucleotides by these HPLC-based methods provides valuable information because G4P/G5P synthesis consumes GDP/GTP and ATP, and because G4P/G5P can function by directly inhibiting GTP biosynthesis (Liu et al., 2015).

G4P/G5P quantification is yet more challenging in plants and algae due to lower concentrations (Takahashi et al., 2004), compartmentation and the high metabolic complexity of these organisms. The first report of G4P and G5P in plants and algae used ion exchange HPLC with UV detection (Takahashi et al., 2004). However, this approach was not widely used-most likely because it requires large amounts of starting material, and a complex extraction procedure. A major step forward for measuring G4P in plants came with the report of a reverse phase HPLC tandem mass spectrometry (MS/MS) based method for G4P quantification that required minimal amounts of starting material and involved a simple solid-phase extraction (SPE) based enrichment step (Ihara et al., 2015). HPLC-MS/MS based quantification allows for greater sensitivity and specificity than HPLC-UV methods. However, a drawback of the Ihara et al. method is that for the calculation of the starting G4P concentration the extract must be split into two aliquots and one aliquot spiked with a known quantity of authentic G4P as an internal standard. This approach corrects for matrix effects, and to some extent for sample loss. However, it is time-consuming, and quantification is vulnerable to inaccuracies caused by the degradation of endogenous G4P before the addition of the internal standard and to differences in treatment between the spiked and un-spiked aliquots. Analyte loss and matrix effects can be overcome using stable-isotope labelled internal standards. Along these lines a method for G4P/G5P quantification in bacteria using metabolically labelled cell extracts and ion exchange HPLC-MS/MS detection was recently described in bacteria (Patacq et al., 2018). Here we build on this previous work and report a new approach for more rapid and accurate G4P quantification in plants and algae using synthesised stable isotope-labelled internal standards. Our approach also allows the simultaneous quantification of GTP, and could be extended to other nucleotides.

## Materials and Methods

### Production and purification of Rel_Seq_ and GppA enzymes

Rel_seq_ (1-385)-6His and 6His-TEV-GppA were expressed in *E. coli*, using plasmids pBAD-Relseq1-385-6His (Mechold et al., 2013) and pET-6his_tev-GppA, which was obtained by cloning the *gppA* ORF between the EcoRI and XhoI restriction sites of the plasmid pEB1188 (Wahl et al., 2011). The proteins were purified by affinity chromatography on cobalt resin as previously described (My et al., 2013)(Supplementary Fig. 1). Final concentrations of Relseq(1-385) and GppA proteins were 4.8 mg.ml^-1^ and 1.4 mg.ml^-1^ respectively, in the purification buffer with 40% glycerol.

### In vitro 13C-G5P and 13C-G4P synthesis

For the synthesis of G5P a reaction containing 2.5 uM Relseq, 2.5 mM ATP, 5 mM 13C-GTP (^13^C_10_ GTP sodium salt; reference 710687, Sigma-Aldrich), 10 mM Tris-Cl pH 8.0, 100 mM NaCl and 15 mM MgCl_2_ was incubated at 37°C for 2 hrs. The reaction mixture was then passed through a spin filter column (Nanosep 10K Omega, Pall Corporation) to remove enzyme before separation of the reaction products by HPLC. For the synthesis of G4P the same reaction as above was performed, and then GppA was added at a concentration of 1.7 uM and the reaction incubated for a further 30 min at 37°C before filtration and separation by HPLC.

### HPLC separation and purification of 13C-G5P and 13C-G4P

An Agilent 1260 Infinity HPLC system with a SAX 5 µm 4.6 × 250 mm Waters Spherisorb analytical column was used to separate samples over 35 min using a gradient of solvent A (KH_2_PO_4_ 50 mM pH 3.4) and solvent B (KH_2_PO_4_ 1 M pH 3.4) at a flow rate of 1 ml min^-1^. The gradient consisted of 5 min, 100% A; 10 min, 70% A; 19 min, 45% A; 20 min, 25% A; 30 min, 0% A; 35 min, 0% A. Nucleotides were identified by absorbance at 254 nm, and selected nucleotides were collected. Collected nucleotides were diluted in formic acid (1M final concentration), loaded onto Oasis WAX SPE cartridges pre-equilibrated with 50 mM ammonium acetate, washed with 50 mM ammonium acetate and then methanol, and then eluted in a mixture of methanol/water/NH_4_OH (20:70:10). Eluates were lyophilised and then resuspended in 5 mM Tris-Cl pH 8.0 before verification by HPLC and HPLC-MS/MS. Dose response curves indicated a linear relationship over a wide range of concentrations for G4P (as observed previously), GTP and their stable-isotope labelled analogues.

### Micororganism and plant growth

#### Escherichia coli

MG1655 (WT) and MG1655 *ΔrelA spoT207*::cat (ppGpp°) (Wahl et al., 2011) were grown in Luria Bertani broth at 37 °C until an OD_600_ of ∼2. MG1655 pALS10 (Svitil et al., 1993) was grown until an OD_600_ of 1.5 and then induced for 1 hr 15 min with 1 mM IPTG. Cells were aliquoted, washed in 1 volume of cold TBS, and pellets frozen at −80°C before nucleotide extraction.

#### Chlamydomonas reinhardtii

cells strain CC-4533 (Li et al., 2016) were grown in TAP medium at 22°C with continuous illumination and agitation at 120 rpm. After 48 hrs when cells had reached a density of around 5 × 10^6^ cells ml^-1^ cells were divided and either cultivated in the light or darkness at 22°C with agitation at 120 rpm. 1.1 × 10^8^ cells were then harvested by centrifugation at 4°C and stored at −80°C until extraction.

#### Arabidopsis thaliana

Col-0 (WT) and OX:RSH3 plants (Sugliani et al., 2016) were grown in soil under a 16-h-light/8-h-dark photoperiod at 18/22°C with 115 µmol m^-2^ s^-1^ PAR fluorescent lighting. After 4 weeks rosette leaves were harvested and ground to a fine powder in liquid N_2_ and then stored at −80°C until extraction.

### GTP and G4P extraction from bacteria, plants and algae

Nucleotides were extracted using a modified version of a previously reported method (Ihara et al., 2015). Samples were mixed with 3 ml of cold 2M formic acid containing stable-isotope labelled nucleotides (125 pmoles 13C-GTP, 12.5 pmoles 13C-G4P) and incubated for 30 minutes on ice. 3 ml of 50 mM ammonium acetate at pH 4.5 was then added, samples were centrifuged at 5000 rpm at 4°C for 10 min and supernatants loaded onto Oasis WAX SPE cartridges pre-equilibrated with 1 ml methanol then 1 ml 50 mM ammonium acetate pH 4.5 at 4°C. After all the sample was loaded, the cartridge was washed with 1 ml 50 mM ammonium acetate pH 4.5 and then 1 ml methanol, and then eluted in 1-2 ml of a mixture of methanol/water/NH_4_OH (20:70:10). Water was added to reduce methanol to 10% and eluates were lyophilised and resuspended in 200 µl water before analysis by HPLC-MS/MS.

### Analysis by HPLC-MS/MS

Extracts were filtered, and analyzed using a Waters Acquity ultra performance liquid chromatograph coupled to a Waters Xevo Triple quadrupole mass spectrometer TQS (UPLC-ESI-MS/MS). 1 µl of extract was separated on a reverse-phase column (Uptisphere C18 UP3HDO, 100 × 2.1 mm, 3 µm particle size; Interchim, France) using a flow rate of 0.3 ml min^-1^ and a binary gradient: (A) 8 mM DMHA with 80 µl AcOH / 500 ml water and (B) acetonitrile, the column temperature was 40 °C. We used the following binary gradient: 0 min, 100 % A; 10 min, 60 % A; 10.5 min, 100 % A; 20 min, 100 % A. Mass spectrometry was conducted in electrospray and Multiple Reaction Monitoring scanning mode (MRM mode), in negative ion mode. Relevant instrumental parameters were set as follows: capillary 1.5 kV (negative mode), source block and desolvation gas temperatures 130 °C and 500 °C, respectively. Nitrogen was used to assist the cone and desolvation (150 L h^-1^ and 800 L h^-1^, respectively), argon was used as the collision gas at a flow of 0.18 ml min^-1^. Multiple reaction conditions and fragment ions selected for quantification are shown in Table 1. Limits of detection (LOD) and quantification (LOQ) were calculated as the nucleotide concentration that could be detected in 150 mg of Arabidopsis tissue with a signal to noise ratio above 3 or 10, respectively (Table 1).

**Table 1.**

| Compound | Precursor | Fragment | Collision energy, mV | LOD, pmol g<sup>-1</sup> | LOQ, pmol g<sup>-1</sup> |
| --- | --- | --- | --- | --- | --- |
| 13C-GTP | 532 | 159 | 35 | 6.3 | 20.7 |
| GTP | 522 | 159 | 35 | 6.3 | 20.7 |
| 13C-G4P | 612 | 159 | 22 | 1.9 | 6.2 |
| G4P | 602 | 159 | 22 | 1.9 | 6.2 |

## Results

Stable-isotope labelled internal standards are considered the gold-standard for the quantification of small metabolites due to their ability to compensate for matrix effects and analyte loss (Stokvis et al., 2005). We therefore set up a method to synthesise stable-isotope labelled G5P and G4P from the widely available stable-isotope labelled GTP. We produced the bacterial enzymes Rel_Seq_, which synthesises G5P, and GppA, which hydrolyses G5P to G4P (see Material and Methods and Supplementary Fig. 1). We then carried out G4P and G5P *in vitro* synthesis reactions as previously described (Corrigan et al., 2016; Mechold et al., 2013) but using totally labelled guanosine-^13^C_10_ 5′-triphosphate (13C-GTP) as a substrate in order to produce 13C labelled G5P and G4P (Fig 1A). We separated the reaction products by ion exchange HPLC and isolated 13C-G5P and 13C-G4P (Figure 1B). HPLC-MS/MS analysis of the isolated 13C-G4P and 13C-G5P showed the expected +10 mass shift for the [M-H]^-^ parent ion compared to the non-labelled compounds, and the same characteristic fragmentation of the parent ion into daughter ions (Fig. 1C).

**Figure 1.**
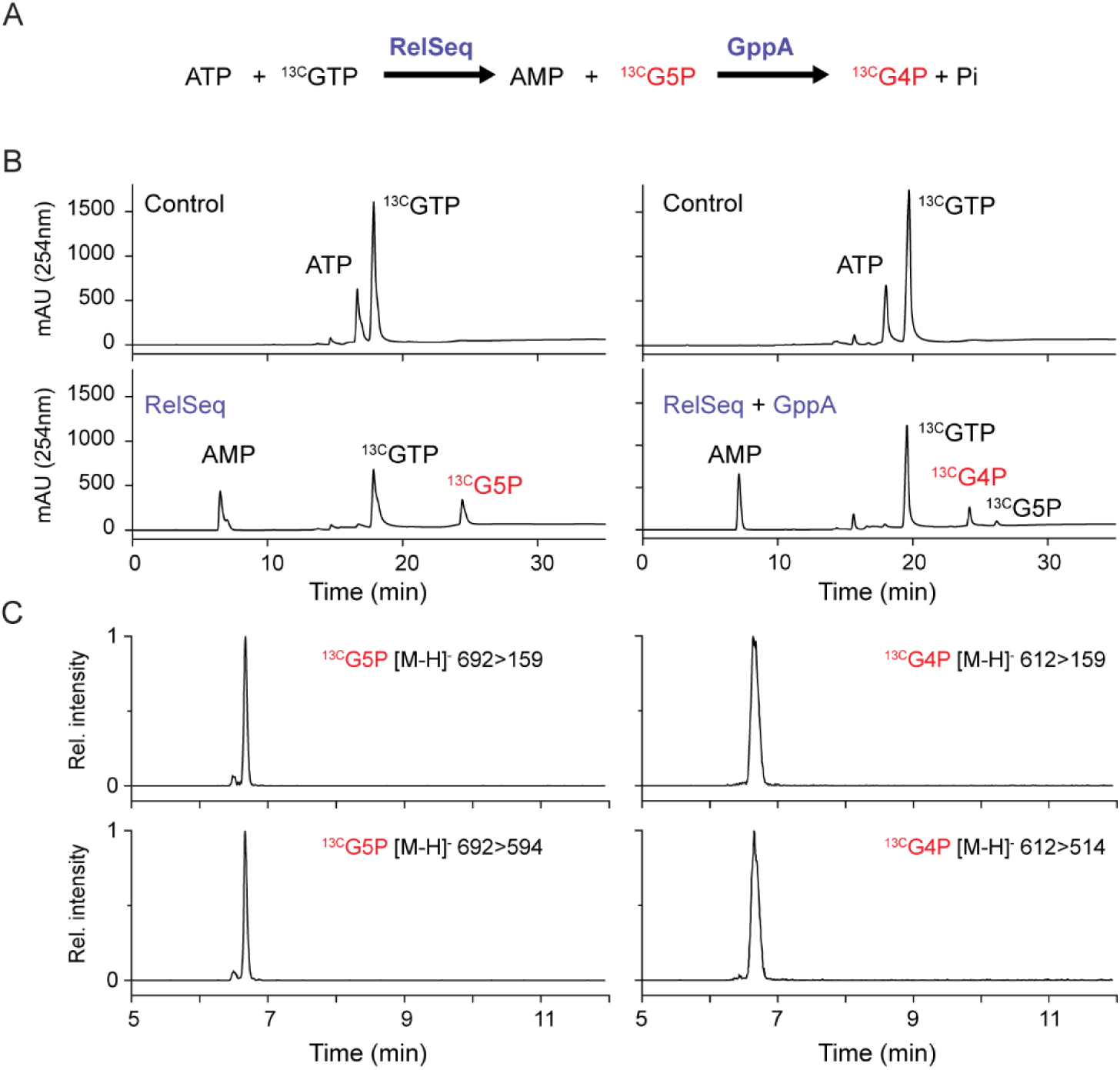
Synthesis and isolation of stable-isotope labelled G4P and G5P. (A) Outline of reaction scheme for the synthesis of isotope-labelled nucleotides. (B) Separation of reaction products by ion-exchange HPLC from control reactions where no enzyme was added, and from reactions where the indicated enzymes were included. (C) Validation of isolated nucleotides by detection of daughter ions following the collision induced ionization of 13C-G5P and 13C-G4P [M-H]- parental ions.

We next confirmed that the stable isotope labelled G4P could be used to quantify endogenous unlabelled G4P levels in bacteria. We adapted the extraction procedure of Ihara et al. (2015) by adding stable-isotope labelled G4P to the extraction buffer before mixing with the biological sample, and by continuing to the SPE enrichment step without splitting each sample (Fig. 2A). This approach allowed us to use the 13C-G4P internal standard signal to account for endogenous G4P loss during the extraction procedure and matrix effects such as ion suppression during MS/MS analysis. Using this method, we calculated endogenous G4P concentrations in wild-type *Escherichia coli*, a G4P/G5P null mutant, and in a strain containing an inducible G4P synthase expressed from plasmid pALS10 (Svitil et al., 1993)(Fig. 2B). Concentrations in the wild-type (24.8 ±2.6 SE pmol OD_600_^-1^) were within the range of steady-state levels previously reported (3-40 pmol per OD) (Hernandez and Bremer, 1991; Maciag et al., 2010; Macvanin et al., 2000; Murray and Bremer, 1996; Slomińska et al., 1999) although higher levels are sometimes detected (Ihara et al., 2015). No G4P was detected in the G4P/G5P null mutant that lacks all G4P/G5P synthetase activity, and three-fold higher levels of G4P were measured in the strain expressing a G4P synthase.

**Figure 2.**
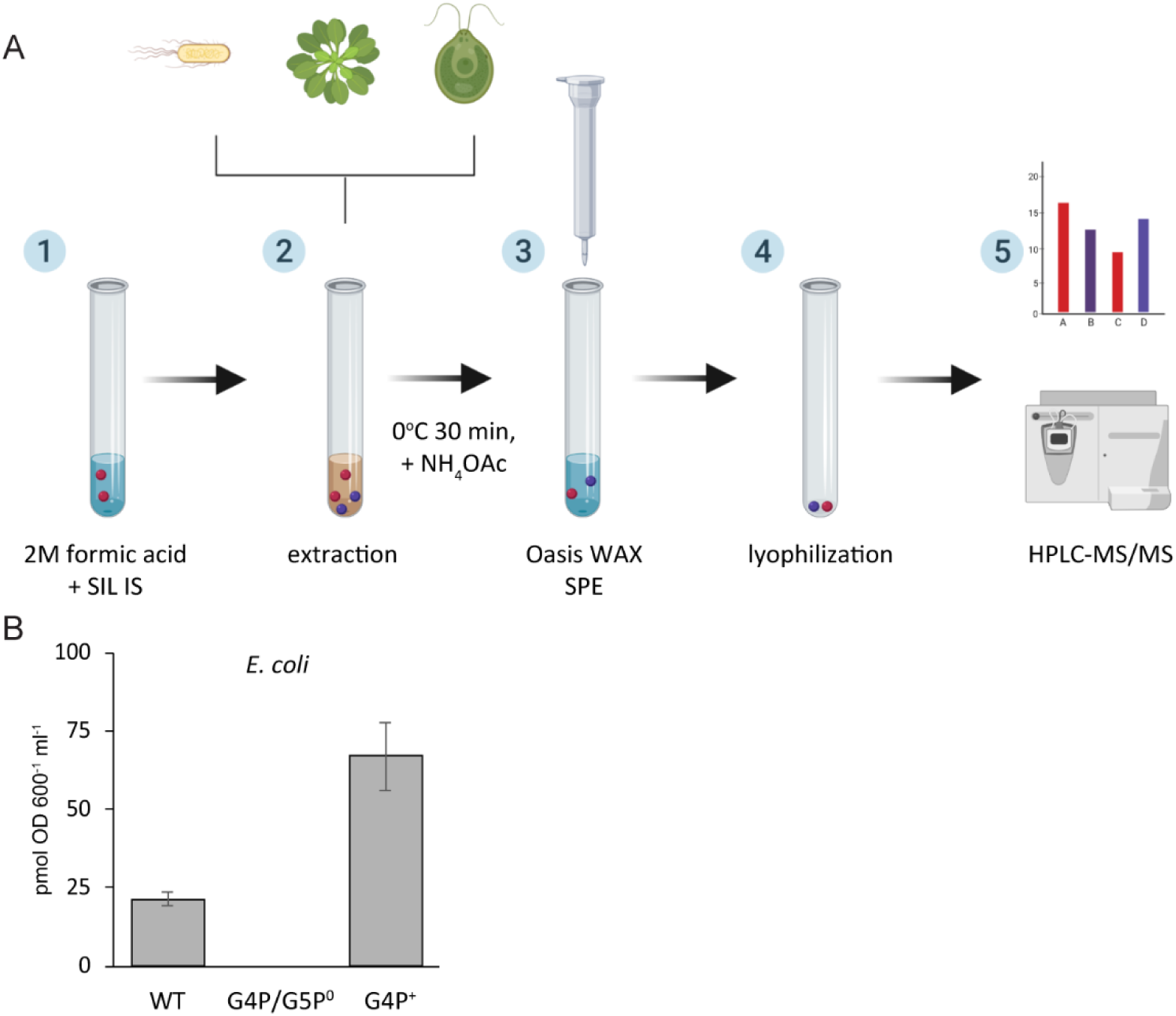
A procedure for the quantification of nucleotides in bacteria, plants and algae. (A) Outline of the extraction procedure with stable-isotope label internal standards. 1, formic acid containing stable-isotope labeled (SIL) internal standards (IS)(red circles) is prepared; 2, biological samples containing unlabeled endogenous nucleotides (violet circles) are added; 3, nucleotides are enriched using an Oasis weak-anion exchange (WAX) solid phase extraction (SPE) cartridge; 4, nucleotides are eluted and lyophilized; and 5, nucleotides are injected into an HPLC-MS/MS system for analysis and quantification. (B) The procedure for the extraction and quantification of G4P was validated on late exponential phase bacterial cells from a wild type strain (WT), a strain lacking G4P and G5P (G4P/G5P°), and a strain over-accumulating G4P (G4P^+^). Means +/- SE are indicated, n= 4 biological replicates.

Once we had established that the method worked in bacteria, we then tested whether stable-isotope labelled internal standards could be used to simultaneously quantify the endogenous levels of GTP and G4P in the more challenging tissues of a plant. Using stable-isotope labelled GTP and G4P internal standards, we were able to quantify endogenous GTP and G4P levels in leaves of wild-type Arabidopsis and plants overexpressing the G4P/G5P synthetase RSH3 (OX:RSH3.1) (Fig. 3A). GTP levels measured in wild-type plants (7.8 ± 0.6 nmol g^-1^) were similar to those previously reported in Arabidopsis (9-16 nmol g^-1^) using ion exchange HPLC with UV detection (Hung et al., 2004; Yang et al., 2015). G4P levels were also similar to those that we previously determined in wild-type (28.7 ± 2.2 pmol g^-1^ versus 21.7 ±3.4 pmol g^-1^) and OX:RSH3.1 plants that overproduce G4P (236.8 ±47.4 pmol g^-1^ versus 147.7 ±7.4 pmol g^-1^) (Sugliani et al., 2016).

**Figure 3.**
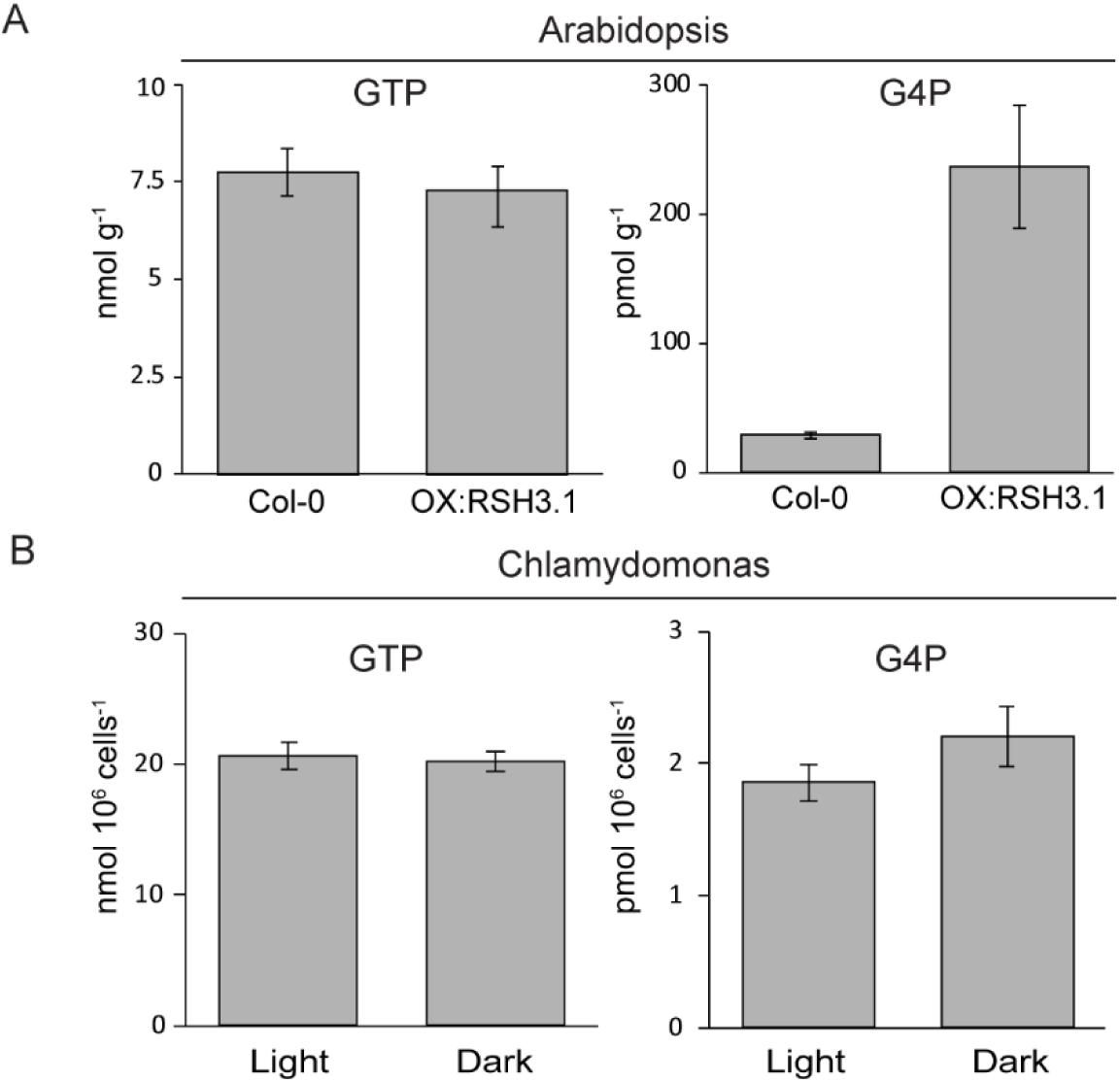
Quantification of GTP and G4P in plants and algae. (A) GTP and G4P were quantified in the leaves of 4 week old Arabidopsis plants using 13-GTP and 13C-G4P internal standards. Means +/- SE are indicated, n= 4 biological replicates. (B) GTP and G4P were quantified in Chlamydomonas cells grown under normal conditions (light) or transferred to darkness for 1 hr (dark) using using 13-GTP and 13C-G4P internal standards. Means +/- SE are indicated, n= 4 biological replicates.

To determine whether this stable-isotope quantification approach could be applied more widely we also simultaneously quantified GTP and G4P in cells of the model green alga *Chlamydomonas reinhardtii* grown under normal conditions or after transfer to darkness for 1 hr (Fig. 3B). Both nucleotides were readily detected, and we observed a small increase in G4P levels in response to dark treatment (*P*=0.1, Wilcoxon test, n=4). The nucleotide concentrations could not be compared to published concentrations because we are not aware of previous reports of GTP or G4P measurements in comparable units in Chlamydomonas.

The 13C-G5P that we produced could be potentially used for quantification, and indeed low amounts of G5P were previously observed in plants (Takahashi et al., 2004). However, in preliminary tests we found G5P detection in plant and algal extracts to be unreliable, suggesting that further optimisation is required. Previously it was also observed that an organic solvent based nucleotide extraction procedure caused the conversion of G5P into G4P, therefore affecting the accuracy of endogenous G4P quantification (Patacq et al., 2018). However, this may be specific to the extraction procedure because we did not observe conversion of 13C-G5P into 13C-G4P following the formic acid-based extraction procedure we describe in this work without addition of biological material.

## Discussion

We describe a stable-isotope internal standard based method for the extraction and quantification of G4P and GTP in plants and algae. Due to the use of stable-isotope internal standards this method is more rapid and accurate than previous methods for measuring G4P and GTP in these organisms, and we hope will accelerate research into the function of G4P signaling in plants, algae and other organisms.

The results generated in this study also raise interesting questions about the relationship between G4P and GTP, and the regulation of G4P biosynthesis in algae. In many bacteria G4P accumulation causes a drop in GTP levels. This can occur via the consumption of GTP, or via the direct inhibition of enzymes in the GTP biosynthesis pathway by G4P (Liu et al., 2015). We observed that OX:RSH3 plants have much higher G4P levels than the wild type control, yet do not have significantly altered GTP content. GTP does not freely exchange between the chloroplast and cytosol (Kusumi and Iba, 2014; Olsen and Keegstra, 1992). Therefore, if we assume that the chloroplast compartment makes a substantial contribution to the cellular GTP pool, then our results indicate that G4P does not inhibit GTP production in the chloroplast in OX:RSH3 plants. This result is somewhat surprising because G4P has been shown to specifically inhibit the chloroplastic guanylate kinase, an enzyme necessary for GTP production in the chloroplast (Nomura et al., 2014).

We also observed only a very small increase in G4P levels in Chlamydomonas following dark treatment. This dark-induced G4P increase is substantially smaller than the 4-6-fold dark-induced increase in G4P that is observed in Arabidopsis and which is dependent on the Ca2+ activated G4P synthase CRSH (Ihara et al., 2015; Ono et al., 2019). A CRSH homologue is also present in *C. reinhardtii*. Our G4P measurements might suggest that there are differences in the strength of CRSH activation in response to darkness between Arabidopsis and *C. reinhardtii*. However, it is also possible that during the preparation of the cells the centrifugation steps, which occur in darkness, cause an increase in G4P in the control sample that masks the increase in the dark-treated cells. Further studies will be required to establish for certain whether dark can induce G4P accumulation.

Finally, we note that our method still has room for improvements. In particular, the HPLC separation approach used here prior to MS/MS detection relies on the ion-pairing reagent DMHA to retain the highly polar nucleotides on a reverse phase C18 chromatography column. Ion-pairing reagents such as DMHA can be problematic for HPLC-MS systems as they cannot be easily removed from columns and plastic tubing, and sometimes require long equilibration times (Yerneni, 2017). Notably, there are recent examples of the successful separation of G4P and other nucleotides from both bacteria and algae using hydrophilic interaction (HILIC) chromatography (Jin et al., 2018; Zborníková et al., 2019).

## Acknowledgements

The authors thank Julie Viala for support in the lab and Yasmine Hassoun for the construction of the GppA expression plasmid. The work was funded by Agence Nationale de la Recherche (ANR-17-CE13-0005) and the IJPB benefits from the support of Saclay Plant Sciences-SPS (ANR-17-EUR-0007).

## Contributions

B.F and E.B conceived the experiments. J.B prepared labelled G4P, J.B and B.F performed nucleotide extractions and S.C and G.M setup the HPLC-MS/MS separation method and quantified nucleotides. J.B, S.C, G. M, E. B and B.F contributed to the analysis and interpretation of the results. B.F and E.B wrote the manuscript. All authors provided critical feedback and helped shape the research, analysis and manuscript.

---

**Figure S1. Purification of Relseq and GppA.** Coomassie Brilliant Blue stained SDS-PAGE gels showing the purification steps for the purification of (**A**) RelSeq and (**B**) GppA. T, total extract; S, soluble extract; FT, flow through; E1-E5, elution fractions; L, size marker ladder; and P, insoluble pellet.

## Notes

#### Summary of Updates

Author affiliations updated. Acknowledgements updated.

