## Supplementary figures and images for "Quantification of guanosine tetraphosphate and other nucleotides in plants and algae using stable isotope-labelled internal standards"

### Figure S1

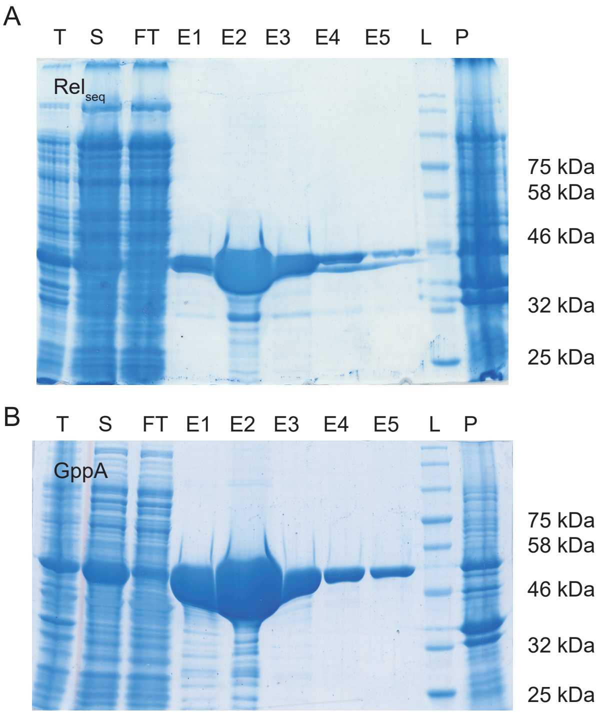
